# Glycolysis Inhibition Regulates Endothelial Junctions by Perturbing Actin and Focal Adhesions

**DOI:** 10.1101/2020.10.22.351148

**Authors:** G. Schwarz, P. Gajwani, J. Rehman, D. Leckband

**Affiliations:** University of Illinois - Urbana-Champaign; University of Illinois at Chicago; University of Illinois at Urbana Champaign

## Abstract

One of the central functions of the endothelium is to maintain a vascular barrier that prevents fluid leakiness and immune cell influx from the circulating blood into the tissue. The barrier integrity of the endothelium is largely controlled by adherens junctions (AJs) and the key AJ molecule VE-cadherin, which maintains cell cohesion via homotypic trans-interaction with VE-cadherin molecules on neighboring endothelial cells. Tension is required to maintain junction homeostasis, but little is known about the role of endothelial metabolism and bioenergetics in regulating junctional forces. Because glycolysis is the main source of ATP generation in endothelial cells, we examined the bioenergetic control of the mechanics of VE-cadherin junctions, by focusing on the glycolysis regulatory enzyme 6-phosphofructo-2-kinase/fructose-2,6-biphosphatase 3 (PFKFB3). Results from traction force imbalance measurements and a VE-cadherin tension sensor revealed that inhibiting PFKFB3 significantly reduced the average junctional tension and the force on VE-cadherin complexes. The decrease in tension was largely due to mechanical changes distal from the cell-cell contacts. Specifically, inhibiting glycolysis perturbs focal adhesions and disrupts actin organization, directly impacting the net force on intercellular contacts. These findings identify a critical role of cellular metabolism for the mechanics and integrity of vascular endothelial junctions, by maintaining global cell mechanics.

**Statement of Significance:** This study examines how forces at intercellular junctions are bioenergetically regulated. Results reveal altered mechanical force generation and transmission due to the breakdown of stress-transmitting fibers during lung injury. These junctions control the barrier function of the vascular endothelium, which requires tight inter cellular adhesions to prevent fluid and macromolecules from passing through the endothelial barrier. We determined how the availability of ATP affects the tension between human endothelial cells, by regulating forces produced remotely from the junctions. These global changes alter both the force at the junctions themselves, and the force transmitted across the entire cell through actin fibers.

## Introduction

Vascular barrier function is a tightly controlled, yet dynamic process that is controlled by adherens junctions, which link neighboring endothelial cells. These junctions are comprised of the surface adhesion molecule vascular endothelial cadherin (VE-Cadherin or Cadherin-5), as well as by catenins, which link the junction complexes intracellularly to the actin cytoskeleton (Cao et al, J Cell Sci, 2019, Le Duc et al., 2010, Tabdili et al., 2011, Buckley et al., 2014). Through this association with actin, the junction complex is maintained under a constant state of tension, which regulates barrier function (Daneshjou N et al, J Cell Bio 2015, Juettner V et al, J Cell Bio 2019, Barry et al., 2014, Huynh et al., 2011). The cells must also respond to external sources of force, such as sheer stress and stretching, which drive remodeling of the junction complexes and associated actin (Salvi et al, Curr Opin Cell Biol, 2018, Tzima et al., 2005, Fang et al., 2019, Gawlak et al., 2014, Ito et al., 2017, Miyake et al., 2006). These active processes are heavily ATP dependent, yet little is known about how endothelial cells derive the energy to maintain their barrier function.

In epithelial cells, this energy need is met by increased glucose oxidation. External force is sensed by E-Cadherin based adherens junctions, which then activate AMPK, leading to increased ATP production through mitochondrial oxidative phosphorylation. This process provides the energy needed for cytoskeletal rearrangements and adherens junction stabilization (Bays et al, Nat Cell Bio, 2017). However, unlike epithelial cells, endothelial cells rely primarily on glycolysis, and not oxidative phosphorylation, for their energy needs (De Bock et al, Cell 2013). In particular, the glycolysis regulatory enzyme 6-phosphofructo-2-kinase/fructose-2,6-biphosphatase 3 (PFKFB3), is a key regulator of endothelial metabolism (De Bock et al, Cell 2013). We thus hypothesized that PFKFB3-driven glycolysis was the source of the ATP required to maintain the integrity of endothelial adherens junctions. The maintenance of tight endothelial junctions is a critical function of the vascular endothelium, because it prevents fluid and protein leak from the blood into the tissue. During pathological processes such as inflammatory lung injury or diabetic retinopathy, this vascular integrity is compromised, and the resulting tissue edema is a key determinant of disease progression (Komarova Y, Circ Res 2017).

In this study, we investigated the impact of PFKFB3 inhibition on VE-cadherin mediated contacts between primary human endothelial cells. Traction force imbalance (TFIB) measurements and the use of a FRET-based VE-cadherin tension sensor (Conway et al, Curr Biol, 2014) revealed mechanical changes at the level of both cell-cell junctions and adherens junction proteins. Using PFK15, which is a pharmacological inhibitor of PFKFB3 (Clem et al, Mol Cancer Ther, 2014), we show that PFK15 treatment significantly decreased both the overall junctional tension and the local force on VE-cadherin molecules. Importantly, these mechanical changes were not confined to local perturbations at cell-cell contacts but reflected global changes in force distributions throughout the cell, including the disruption of focal adhesions and loss of cytoskeletal coherence. These findings reveal the importance of glycolysis in maintaining global cell mechanics, which in turn regulates barrier homeostasis in endothelial cells.

## MATERIALS AND METHODS

### MATERIALS

#### Cell lines, adhesive protein and inhibitor

Human Lung Microvascular Endothelial Cells (HLMVECs) obtained from Cell Applications Inc. were maintained in growth medium EGM-2MV (Lonza) medium supplemented with 10% (v/v) fetal bovine serum (FBS, Sigma – F0926-500ML) and 1% (v/v) penicillin– streptomycin (Corning Cell Grow, Manassas, VA). Glycolysis was inhibited, by incubating cells with 10 μM PFK15 (Cayman Chemical - #17689) for 40 min, prior to measurements. Cells adhered to substrates coated with fibronectin (Sigma SIGMA-ALDRICH INC. - FC010-5MG) diluted 10× (0.1mg/mL) with phosphate buffered saline (PBS) (Corming, #21-040-CV). HEK293T and HEK293AD cells were cultured in DMEM (Corning) supplemented with 10% FBS and 1% PenStrep (Gibco).

#### Preparation of polyacrylamide gels

The 35 mm diameter glass bottom dishes with13 mm wells (Cell E&G - #GBD00001-200) were treated with 0.1M NaOH in order to make the glass hydrophilic. Then the glass was treated with APTMS (Sigma), washed twice, and treated with (v/v) 0.5% gluteraldyhyde (Sigma) to covalently link the polyacrylamide gels to the glass. The Young’s modulus of the polyacrylamide gels used for the experiments was 40 kPa. Gels were prepared with deionized (DI) water, 8% acrylamide, 0.48% bis acrylamide, and 2μL of 0.2μm red fluorescent beads (Life Technologies Corporation - F8810) (Engler et al., 2004). The latter beads are used as fiducial markers for traction force measurements. Crosslinking agent ammonium persulfate (10% w/v APS, Sigma Aldrich) and catalyst N,N,N′,N′-tetramethylethylenediamine (0.5 μL - TEMED, Sigma Aldrich) was used to initiate the reaction. Next, 20 μL of the gel mixture was pipetted onto the center of each glass bottom dish and covered with a 12 mm diameter glass cover slip (EMS Acquisition Corp / Electron Microscopy Sciences - 72231-01). After curing gels for 30 minutes at 37°C, the gels were submerged in DI water to keep the gels hydrated and to remove any unreacted chemicals. Gels were stored at 40°C for up to 2 weeks.

#### Preparation of PDMS stamps

Polydimethylsiloxane (PDMS) stamps were created to microcontact print fibronectin in patterns of tangential 1600 μm^2^ circles on 40 kPa gels. To create the PDMS stamps, 10:1 Sylgard 184 Silicone Elastomer Base, part B, (Sylgard^tn^ 4019862) Sylgard 184 Silicone Elastomer Curing Agent, part A, respectively. The mixture was poured over a negative master and put into a decanter for 1 hour, to remove bubbles. Then the mixture was cured for 15 minutes in a 108° C oven. The PDMS was peeled from the master and cut into individual stamps (Qin et al., 2010)

#### Viral Infections

##### Plasmid Constructs

The FRET-based VE-Cadherin tension sensor (VE-CadTS: (Addgene Plasmid# 45848) was used to measure tension on VE-cadherin bonds. For expression in endothelial cells, both constructs were cloned into the PLJM1-EGFP lentiviral vector (Addgene, Plasmid #19319). The inserts were amplified by PCR, using the Q5 Polymerase (New England Biolabs) and ligated into the digested PLJM1 vector at AgeI and EcoRI restriction sites. Cloning resulted in the removal of EGFP from the vector backbone. Successful cloning was confirmed by Sanger sequencing. Additional confirmation was obtained by observing the correct reporter localization at cell-cell junctions, as visualized under a confocal fluorescent microscope.

##### Virus Production

Lentivirus was produced to express VE-CadTS and VE-CadTL in HLMVECs. To generate the virus, VSV-G (lentiviral envelope expressing vector, Addgene, Plasmid# 12259), psPAX2 (viral packaging vector, Addgene, Plasmid# 12260) and either PLJM1-VE-CadTS or PLJM1-VE-CadTL were co-transfected into HEK293T cells using JetPrime (Polyplus) according to the manufacturer’s protocol. The virus-containing supernatant was harvested 48 hours after transfection, and the viral particles were precipitated using Lenti-X concentrator (Takara Bio) following the manufacturer’s protocol. Low passage HLMVECs were transduced with concentrated virus, along with 10μg/mL Polybrene (Millipore Sigma). Expression was observed 2-7 days following infection.

Adenovirus was used to express VE-Cadherin-GFP in HLMVECs (Shaw et al, 2001). The virus was amplified in HEK293AD cells, by transducing the cells and collecting the virus containing supernatant, once the cytopathic effect was complete. The viral particles were then purified using the Adeno-X Maxi Purification Kit (Takara Bio) as per the manufacturer’s instructions. Low passage HLMVECs were transduced with the virus, and expression was observed 2-3 days following infection.

#### Viral infections for GFP-VE-cadherin and VE-CadTS expression in HLMVECs

Viral infections were performed in accordance with established BSL2 protocols. HLMVECs were cultured in Human Microvascular Endothelial Cell Medium (HLMVEC Cell Applications) with 10% FBS and 1% penicillin-streptomycin. Cells grew to 60% confluence in 6-well plates before treating with the viral infections. The VE-CadTS and polybrene were added to the cells for the lentiviral infection whereas just the VE-cadherin GFP virus was added for the adenoviral infection. Cells were incubated with virus at 37°C under 5% CO_2_ for 12-18 hours, before the medium was changed to virus-free medium. Cells were then cultured for 2 additional days, to allow for fluorescent protein expression.

### METHODS

#### Micro contact printing fibronectin patterns on polyacrylamide gels

PDMS stamps were submerged in ethanol and sonicated for 5 minutes and plasma treated (Harrick Plasma - PDC-32G) to make the stamps hydrophilic. Fibronectin (0.1mg/mL) was pipetted onto the stamps and incubated for 1 hour, to allow fibronectin to adsorb on the stamps. Fibronectin was immobilized on Sulfo-SANPAH-activated (25mg/μl), 40 kPa gels by aspirating the PBS from the stamp and then stamping the adsorbed protein onto the activated gels. Gels were incubated at 37° C before the stamps were removed. The gels with printed fibronectin patterns were then submerged in PBS and stored overnight at 4° C.

#### Traction force microscopy

Traction force microscopy (TFM) measurements used polyacrylamide hydrogels with Young’s moduli of 40 kPa. Fibronectin (0.1mg/mL) was immobilized on Sulfo-SANPAH-activated gels, by incubation with 0.1 mg/ml fibronectin for 4 hr at 37°C (Sigma SIGMA-ALDRICH INC. - FC010-5MG). After incubation, gels were washed with 1x phosphate buffered saline (PBS). HLMVECs were seeded on the coated gels and incubated overnight at 37°C under 5% CO_2_ to allow cells to reach mechanical equilibrium. HLMVECs were treated with PFK15 for 40 minutes. Control cells were untreated. The root mean squared (RMS) traction stress was determined, by comparing the position of the beads beneath the cells, before and after they were lysed. Traction force heat maps were calculated using a custom Matlab program (Butler et al. 2001), provided by Ning Wang (University of Illinois at Urbana-Champaign).

#### Traction force imbalance (TFIMB) measurements

HLMVECs expressing the GFP-tagged VE-Cadherin were seeded at ~ 20,000 cells per gel and incubated at 37°C under 5% CO_2_ overnight, to allow cells to reach mechanical equilibrium. Cells adhered to the stamped fibronectin patterns. We determined the traction forces exerted by the cells (Butler et al. 2001). To determine the net force orthogonal to the cell-cell junctions, we used a custom Traction Force IMBalance (TFIMB) program, which was based on a prior report (Maruthamuthu et al., 2011). We calculated the unconstrained traction forces because the boundary conditions used to calculate constrained tractions force maps are incompatible with the TFIMB calculations. The junction tension (force/length) was calculated by dividing the average force orthogonal to the cell-cell contact by the junction length, which was visualized by imaging GFP-VE-cadherin.

#### Immunofluorescent imaging

After cells were treated with PFK15 for 40 min, they were fixed with 2% paraformaldehyde for 15 min at 37°C under 5% CO_2_. Control cells were not treated with inhibitor. Post fixation, cells were washed 3x with PBS for 5 minutes each on a slow rocker. Cells were permeabilized with (v/v) 0.1% 100x Triton mix (Sigma - T8787-100ML), washed 3x for 3 minutes with PBS on a slow rocker, and then incubated with a blocking buffer containing 1%(w/v) Bovine Serum Albumin (BSA), for 20 minutes. Then, the cells were treated with the primary antibodies and phalloidin for 1 hour. Focal adhesions were visualized by immunostaining paxillin with primary mouse anti-paxillin antibody (1:200 BD Transduction Laboratories – 3108546-612405) and secondary antibody goat anti-mouse 488 (1:200 Invitrogen - 41819A). VE-cadherin was immunostained with rabbit anti-VE-Cadherin (1:200 Cell signaling Technologies - D87F2) followed by secondary, goat anti-rabbit Cy5 (1:200 Invitrogen - 1843847). F-actin was stained with phalloidin 488 (1:200 Invitrogen - A 12379). After labeling with primary antibodies, the cells were washed five times at 5 min per wash, and then treated with the secondary antibodies. Finally, the cells were washed 5 times at 5 min per wash, with the last wash containing DAPI at a (v/v) 1,000x dilution. Cells were then treated with mounting medium (Invitrogen - 2147630). Images were acquired with a Zeiss Axiovert 200M wide view microscope at 40x with an oil immersion objective, using Carl Zeiss™ Immersol™ Immersion Oil (Fisher Scientific - 12-624-66A). VE-cadherin was imaged by illumination at 647 nm laser, and the paxillin and F-actin fluorescent labels were excited at 488 nm.

#### FRET measurements of junction tension

FRET images of junctions between HLMVECs expressing the VE-cad-TS were imaged at 40x with a Zeiss Axiovert 200M wideview microscope. Regions of interest were imaged at each junction with a CFP (filter set 47, #000000-1196-682, excitation: BP 436/20, emission: 480/40) filter and a FRET (CYP/YFP filter set 48, 000000-1196-684, excitation: BP 436/20, emission: 535/30) base pair filter. Intensities at the junction with the CFP filter, FRET filter, and background were quantified in imageJ. The background of each was subtracted for both filters in ImageJ, by measuring the background-subtracted intensity at the junctions. FRET/CFP ratios were calculated using ImageJ. As increased FRET indicates a decrease in FRET, the inverse ratio, CFP/FRET was graphed. Thus, an increase in CFP/FRET indicates an increase in tension.

For the measurements, ~ 20,000 cells were seeded onto each microcontact-printed, 40 kPa gel, prepared as described above. Cells were incubated overnight at 37°C under 5% CO_2_, in order for cell doublets on the patterns to reach mechanical equilibrium. Before imaging, the cells were treated with the inhibitor, as described above. Control cells were not treated.

#### FRAP measurements of cadherin mobility

We conducted fluorescence recovery after photobleaching (FRAP) measurements of VE-cadherin at junctions between HLMVECs in a confluent monolayer, using a Confocal – Zeiss Light Scanning Microscope 880. GFP-VE-cadherin was expressed in HLMVECs following adenovirus infection, as described above. The GFP-VE-cadherin fluorescence in a region of interest of 17 μm^2^ was photobleached for 500 ms at the maximum laser intensity of ~9.3 W/cm^2^ and then allowed to recover with the laser at a reduced intensity of ~1.04 W/cm^2^ for 10 minutes, with a recovery area of 29 μm^2^. Intensities were measured continuously and recorded every 500ms during the measurements at 20x magnification and a wavelength of 488 nm. The intensities at the different time points were normalized relative to the pre-bleach intensity.

HLMVEC monolayers at ~85% confluence were cultured on fibronectin-coated, 40 kPa gels, as described above. Cells were incubated overnight at 37°C and under 5% CO_2_. Prior to measurements, cells were treated with the inhibitor, as described above. Control cells were not treated.

## Results

### Cell adhesion on microcontact printed fibronectin patterns on 40kPa poly (acrylamide) gels

Measurements of the force per unit length (tension) at junctions between human lung microvascular endothelial cells (HLMVECs) were carried out with cells on arrays of microcontact-printed fibronectin on polyacrylamide (PAA) gels. The stamped fibronectin pattern of two, tangential 1600 μm^2^ circles is shown in Figure 1a, and a representative fluorescence image of micropatterned arrays of fluorescently labeled fibronectin is shown in Figure 1b.

**Figure 1.**
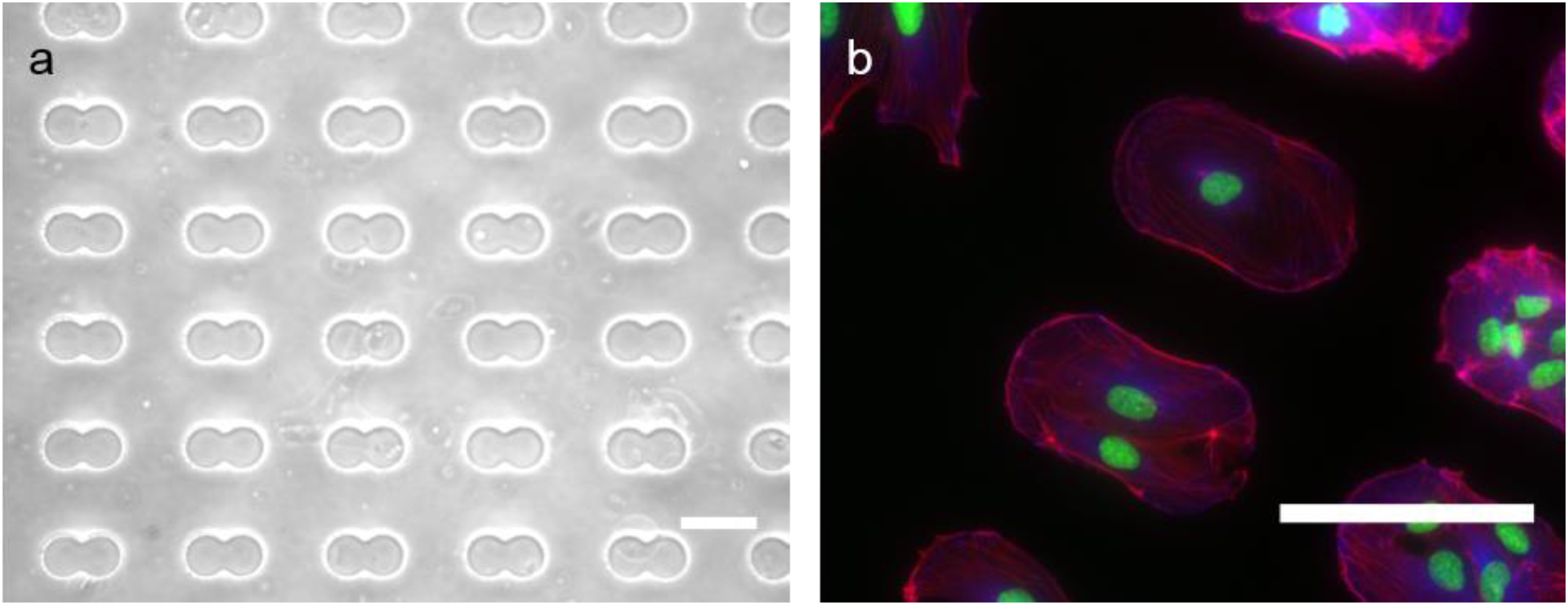
Cell doublets on micro contact printed fibronectin patterns on polyacrylamide gels : a) PDMS stamp pattern of tangential 1,600 μ^2^ circles. Scale bar = 90 μm. b) Cell doublets on fibronectin patterns printed on 40kPa gels. Cells were immunostained for paxillin (blue), actin (red), and DAPI (green). Cells were imaged at 40x magnification with a scanning confocal microscope. Scale bar = 45 μm.

Cells adhered to the patterns (Fig. 1b), but the shapes of cells on these patterns were variable, such that the junction lengths and junction orientations relative to the long axis of the doublet differed. We determined the junction tension from the average traction stress exerted by each cell, but cell shapes can affect the traction generation. We therefore conducted sufficient traction force imbalance (TFIMB) measurements to ensure that we accounted for the variability in cell shapes and could thus determine the tension differences between control cells and those with inhibition of glycolysis via the PFKFB3 inhibitor PFK15.

### Inhibiting glycolysis reduces force on VE-cadherin adhesions

Treatment with PFK15 correlated with a decrease in the force on VE-cadherin bonds. Studies compared the CFP/FRET ratio of the VE-cadTS at junctions between patterned HLMVECs, in the absence and presence of PFK15. Figure 2 shows the fluorescence images of untreated cells (Fig. 2a) and of cells 40 min after PFK15 treatment (Fig. 2b). As determined from the normalized CFP/FRET ratios (Fig. 2c), glycolysis inhibition reduced the normalized CFP/FRET ratio from 0.97 ± 0.06 to 0.65 ± 0.03, indicative of a loss in force on VE-cadherin complexes. This difference was statistically significant (p < 0.001, NC = 29, NI= 34).

**Figure 2:**
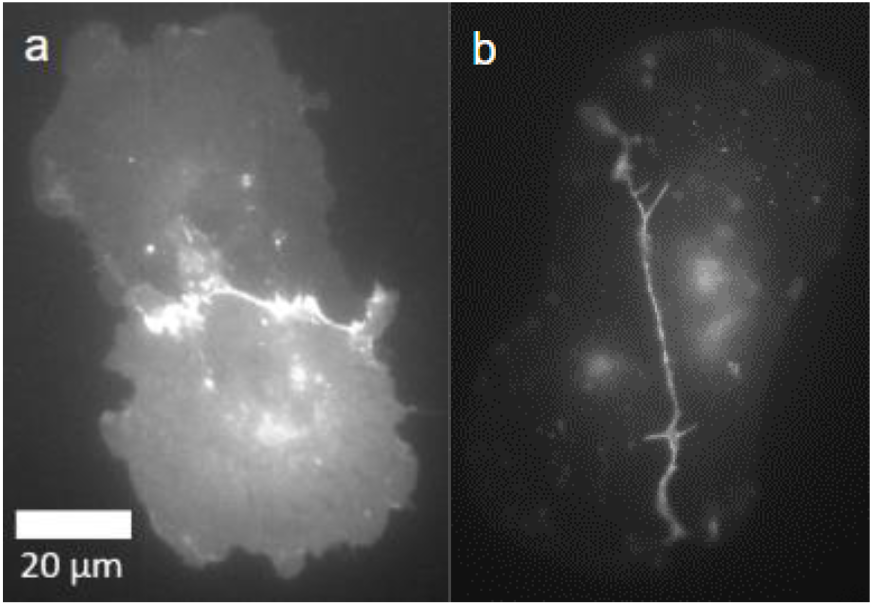
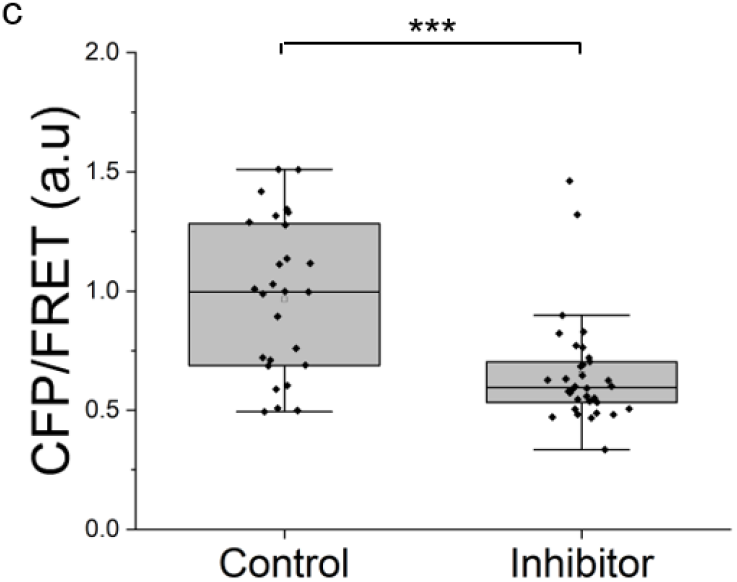
PFK15 treatment reduces the tension on VE-cadherin mediated junctions between HLMVEC pairs. a) Fluorescence images of the VE-cadTS at the junctions between a) control and b) PFK15 treated HLMVECs expressing VE-cadTS (images obtained with the YFP channel were enhanced for presentation purposes only). c) CFP/FRET ratio measured in the region of interest at cell-cell junctions. Average CFP/FRET is significantly lower in the PFK15 treated cells than in the control (***p < 0.001, N_C_ = 29, N_I_ = 34).

We also observed morphological differences. Junctions between untreated cells were broad and typical of the overlapping contacts between endothelial cells. Although the width of the junction between the PFK15 treated cells in Fig. 2b appear narrower and straighter, this apparent difference was not observed uniformly across cells. Fig. 2b also shows some membrane invaginations at junctions between PFK15 treated cells that were observed more frequently than with treated cells.

### PFK15 treatment alters cell tractions and tension at cell-cell junctions

To determine the effects of glycolysis inhibition on the average junction tension, which depends on global cell mechanics as well as on local changes at intercellular adhesions, we performed Traction Force IMBalance measurements (TFIMB). Measurements compared the tension at the junctions, with and without PFK15 treatment. Figures 3a&d are DIC images of HLMVEC doublets on a 40 kPa gel, at 40 min without (Fig. 3a) and with (Fig. 3d) PFK15 treatment. Figures 3b&e compare the traction stresses exerted by untreated and inhibitor-treated cells, respectively. Comparison of Figs. 3b&e shows that PFK15 treatment decreases the traction stresses at the basal plane, relative to untreated controls (Fig. 3b). This is apparent in the red (high stress) to blue (low stress) shifts in the heat maps.

**Figure 3:**
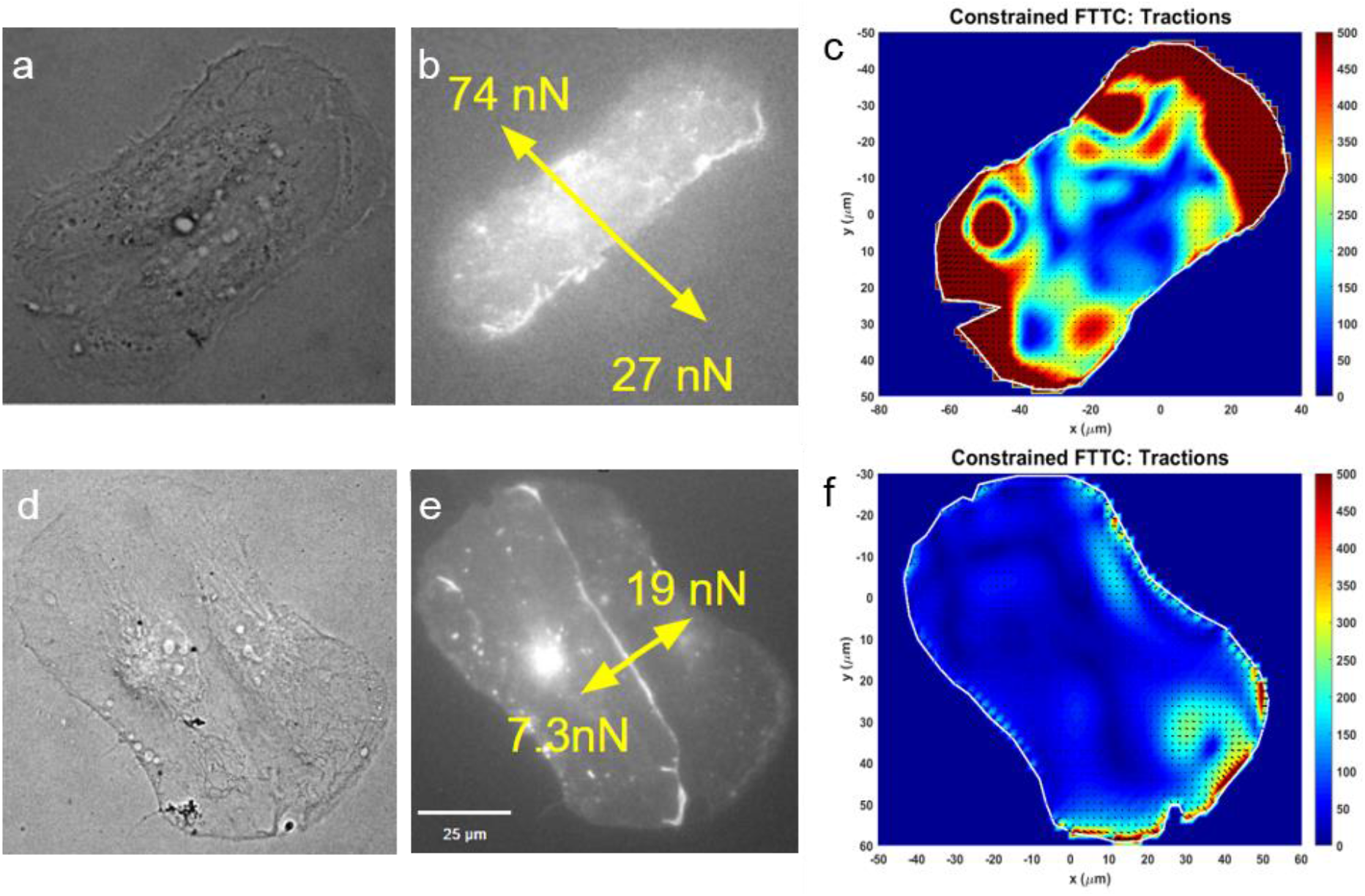
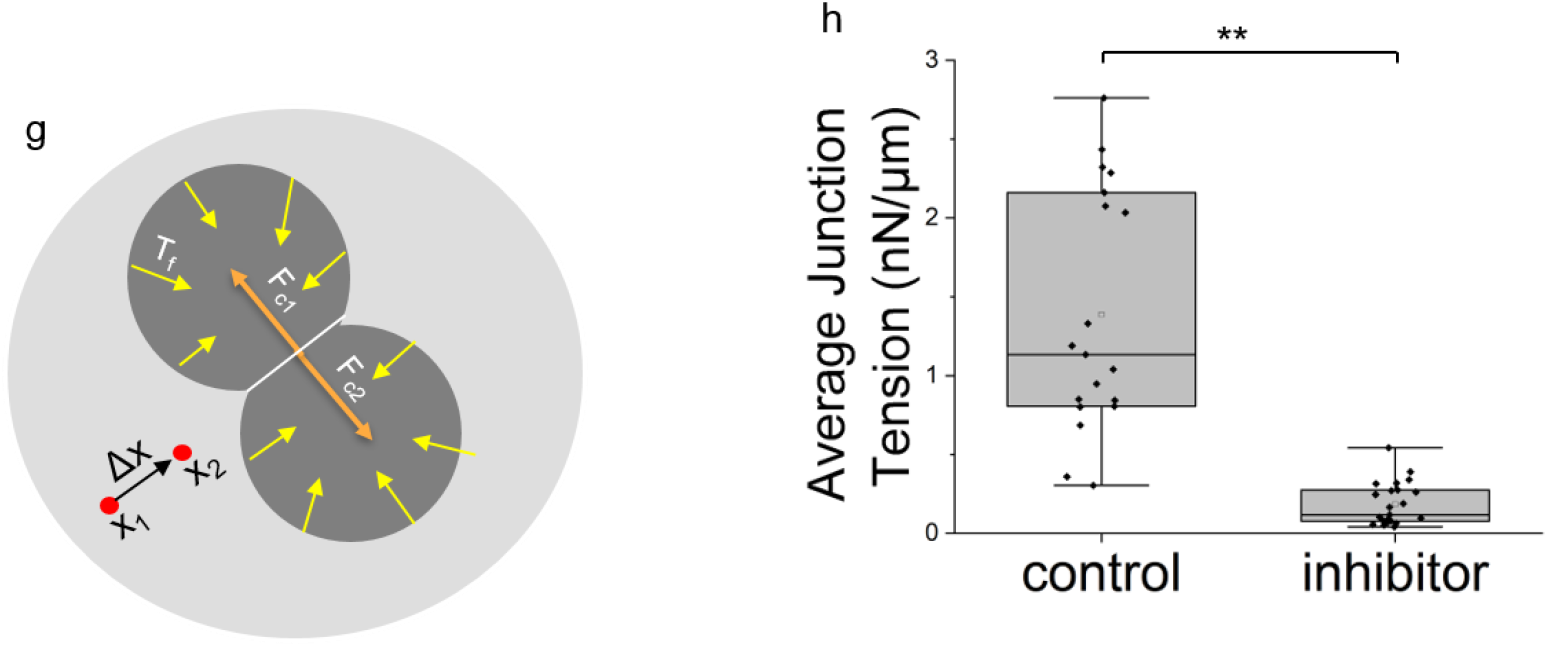
Traction Force Imbalance (TFIMB) measurements of HLMVEC pairs. Substrates were 40 kPa gels. DIC Images of untreated (a) and PFK15 treated (c) cell pairs on fibronectin patterns. Fluorescence image of control (b) and PFK15 treated (e) cells expressing GFP-VE-Cadherin and the determined forces exerted on the junction by each cell. Heat maps of traction stresses exerted by (c) control and (f) PFK15 treated HLMVEC doublets seeded on stamped fibronectin patterns of tangential circles. (g) Force diagram of HLMVEC doublets on a substrate showing the traction force vectors relative to the axis of the junction. Tr is the traction force, F_c1_ and F_c2_ are the forces exerted by cell 1 and cell 2, respectively, and Δx is the length of the junction. (h) Average junction tension of control and PFK15 treated cells (p < 0.01, N_C_ = 20, N_I_ = 24).

Figures 3c&f display the net force on the junction produced by each cell in the doublet. Junctions were visualized with GFP-VE-cadherin. Figure 3g illustrates the force diagram underlying the TFIMB measurements. Here, displacements of the beads in the gels are split into their x and y components, which define the x and y components of the traction stresses. The latter are used to determine the net force orthogonal to the junction. The difference in traction stresses is balanced by the net force on the junction (Maruthamuthu et al., 2011). The average junction tension (Fig. 3g) is the net force divided by the junction length, which is visualized with GFP-VE-cadherin.

In the absence of PFK15 (Fig. 3c), force balance analysis indicated an average net positive force on the junctions. As shown in Fig. 3f, PFK15 treatment decreases the magnitudes of the net force exerted by each cell, with a corresponding decrease in the junction tension, compared to untreated cells (Fig. 3c). Quantitatively, the average tension decreased from 1.7 ± 0.4 to 0.3 ± 0.1 nN/μm. This statistically significant difference (p < 0.01, NC = 20, N_I_ = 14) is qualitatively consistent with the VE-cadherin tension sensor results (Fig. 2). This result reveals that PFK15-dependent changes in global cell mechanics reduces the inter-endothelial tension.

### PFK15 treatment alters the magnitudes and orientations of traction stress vectors

PFK15 disrupted the magnitude and orientational coherence of the individual traction vectors exerted by the cells. The direction and magnitude of the forces determine the junction tension, so we investigated the distribution of vector orientations, with and without PFK15 treatment. Figures 4a&b show wind rose plots of the angles of the traction stresses relative to the junction normal, in the absence and presence of inhibitor, respectively. In untreated cells (Fig. 4a), the absolute values of the angles between the traction force vectors and the junction normal are primarily between 0° and 35° such that the traction vectors are largely oriented perpendicular to the junctions. By contrast, the stress vectors exerted by PFK15 treated cells (Fig 4b) appear to be more randomly distributed.

**Figure 4:**
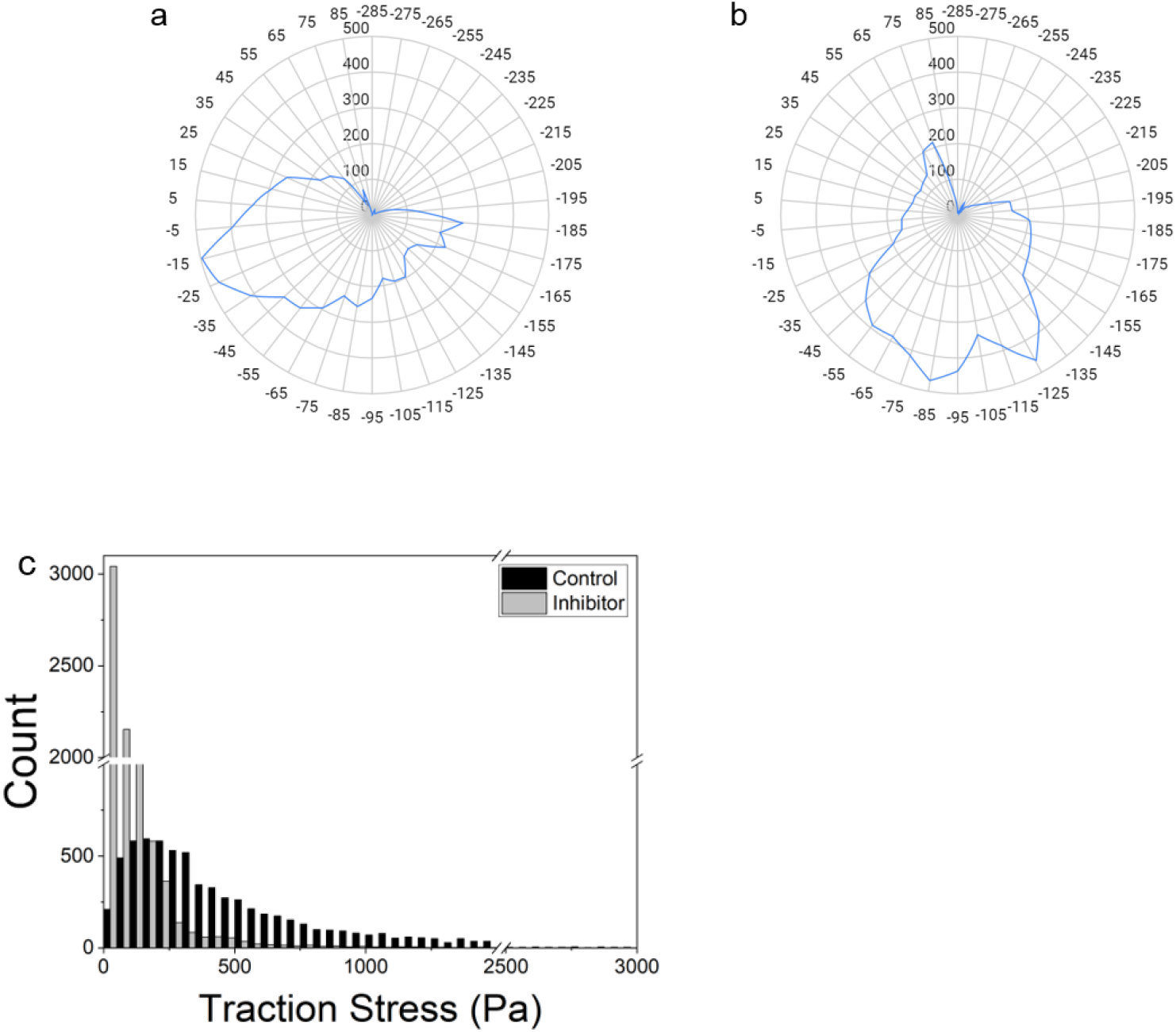
Traction stress distribution and orientation. Wind rose plots of the angles of traction force vectors relative to the junction normal in (a) control and (b) treated HLMVEC doublets. (c) Histogram of the magnitudes of measured traction stresses exerted by cells doublets on printed fibronectin patterns on 40 kPa gels. Black bars indicate control cells (N_C_ = 20) and gray bars were obtained with inhibitor treated cells (N_I_ = 24).

Inhibiting glycolysis also altered the distribution of the magnitudes of the traction stresses. This is shown in the histograms in Figure 4c. In the untreated cells, the most probable traction stress is ~200Pa, and there is a broad, long tail. With PFK15 treated cells, the most probable traction stress decreased to ~50Pa, the distribution is narrower, and the tail is substantially shorter. Taken together, these results show that inhibiting glycolysis disrupts both the relatively uniform orientation traction stresses normal to the junctions (Figs. 4a,b) and shifts the distribution of traction stresses to smaller values (Fig. 4c).

We confirmed the impact of PFK15 treatment on global cell mechanics, independent of cell-cell junctions, by analyzing traction stresses of individual cells, using Traction Force Microscopy (TFM). Figure S1 compares traction stress heat maps of individual HLMVECs on 40 kPa gels without (Fig. S1a) and after (Fig. S1b) PFK15 treatment. In untreated cells, the force vectors are oriented radially inward (Fig. S1a) and actin fibers span the width of the cell, as reported previously (Cai et al., 2010). PFK15 treatment resulted in more randomly oriented stress vectors (Fig. S1b). There was also a drop in the magnitude of the root mean square (RMS) traction stress exerted by individual cells from 210 Pa ± 20 to 100 ± 10 Pa (p < 0.001, N_C_ = 28, N_I_ = 29).

### Glycolysis inhibition disrupts actin networks and focal adhesions

To investigate subcellular changes underlying the decreased magnitude and orientational coherence of the traction stress vectors, following PFK15 treatment, we imaged F-actin and focal adhesions. DAPI staining visualized the nuclei of the individual cells, and GFP-VE-cadherin visualized the cell-cell junctions. In untreated cells, paxillin staining of focal adhesions at the basal plane was distinct and punctate (Fig. 5a), with larger focal adhesions at the cell perimeter. Paxillin was also depleted in the region adjacent to cell-cell junctions. Actin stress fibers were prominent at the basal plane, and crisp F-actin fibers spanned the length of the cell (Fig. 5b). Dense cortical F-actin staining was also apparent parallel to the cell-cell junctions. Vector maps of the orientations of actin fibers revealed the relatively uniform fiber alignment, relative to junctions, in untreated cells (Fig. 5c).

**Figure 5:**
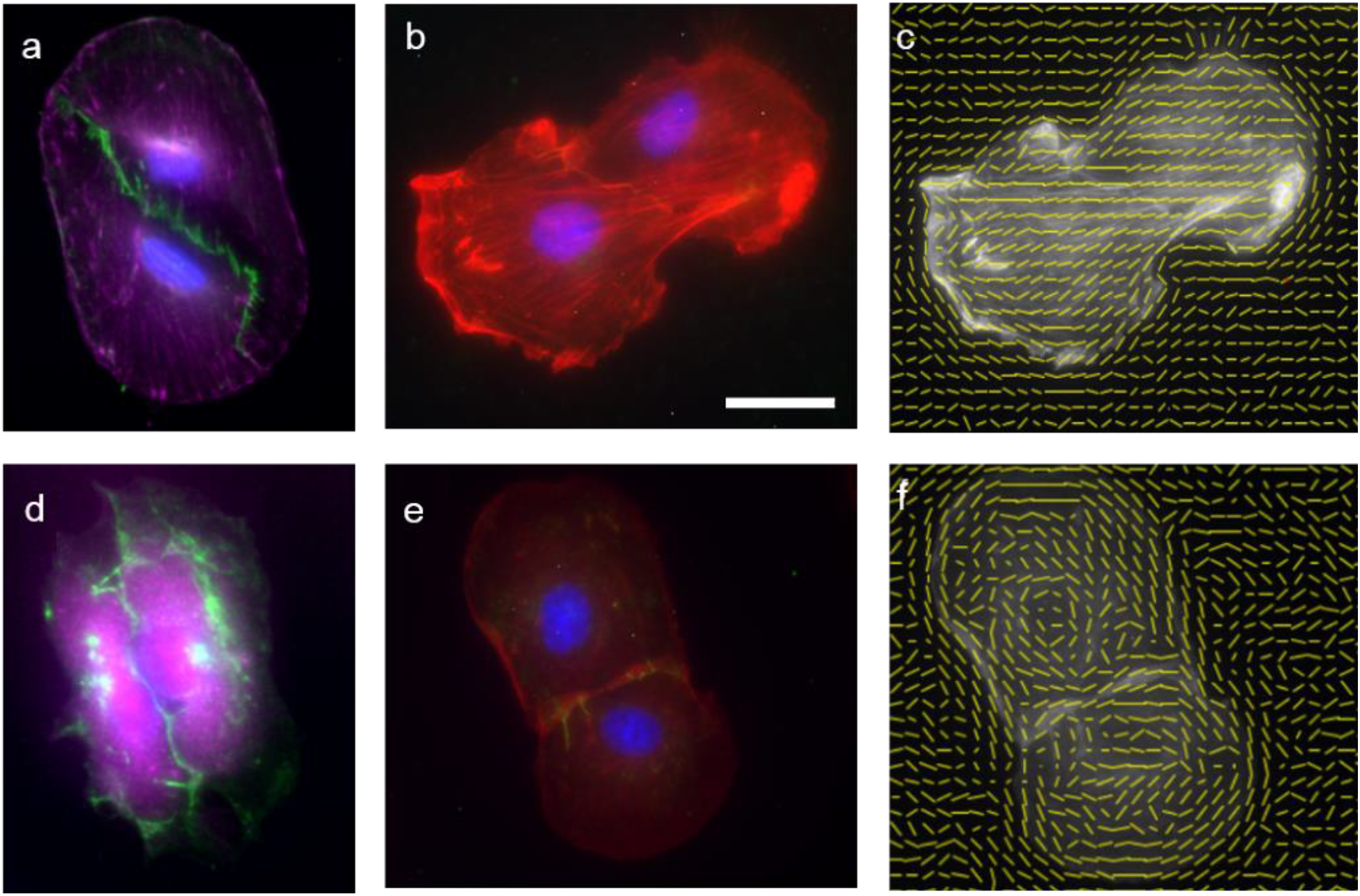
Inhibiting glycolysis disrupts focal adhesions and F-actin: Merged immunofluorescence images of paxillin (purple) and VE-cadherin (green) in a) untreated and d) PFK15 treated HLMVEC doubles. Merged immunofluorescence images of actin (purple) and VE-cadherin (green) in b) untreated and e) PFK15 treated HLMVEC doublets. Nuclei are stained with DAPI. c/f) Vector maps showing the directionality of the actin fibers in c) untreated and f) PFK15 treated HLMVECs. The vector maps correspond to cells in panels b & e. In all images, cells were on micro contact printed fibronectin on 40 kPa gels. Scale bar = 25 μm.

PFK15 treatment disrupted both focal adhesions and F-actin. Paxillin staining was more diffuse (Fig. 5d), and puncta at the basal plane were fewer and smaller. F-actin was also more diffuse throughout the cell (Fig. 5e), and there were fewer, well-defined actin fibers. The vector maps (Fig. 5d) of actin fibers in treated cells also revealed more random fiber orientations.

### Glycolysis inhibition disrupts F-actin fibers and reduces fiber anisotropy

To visualize the change in the actin fibers between the PFK15 treated and untreated cells, we used ImageJ’s “plot profile” tool to generate intensity scans associated with the F-actin across the cell. The scan direction was orthogonal to the majority of visible actin fibers. We examined the phalloidin intensity profiles (Fig. S2 a&c) of both PFK15 treated and untreated cells across the nucleus. In scans of untreated cells, there are distinct intensity peaks (Fig S2b) and valleys associated with the fibers. In PFK15 treated cells, the fibers aren’t as distinct: scans contain fewer of the larger peaks and valleys (Fig. S2d). The large drop in actin intensity across the nuclei also revealed a significant reduction in perinuclear actin.

To characterize the difference in actin fiber alignment in treated versus untreated cells, we used the ImageJ plugin “Fibriltool” (Boudaoud et al., 2014), which calculates the fiber anisotropy and a directionality vector within a region of interest (ROI). Here the ROI is the entire cell. Figure S3 shows the cell doublets and the ROI (yellow outline), as well as the magnitude and direction of the of the calculated actin orientational anisotropy (red line). In untreated cells, the fiber orientation is anisotropic, with the majority of fibers directed along the long axis of the cell doublets and nearly perpendicular to the junctions. The calculated anisotropy value for the untreated cells is significantly larger than that of PFK15 treated cells, as indicated by the relative lengths of the red lines. The anisotropy number for actin in PFK15 treated cells decreased from 0.24 ± .09 (SEM) to 0.013 ± .012 (SEM) (Fig. S3; N_control_ = 46, N_inhibiter_ = 33, p < 0.005).

### Inhibiting glycolysis reduces the diffusivity and mobile fraction of junctional VE-cadherin

Actin disruption would also affect local processes at inter cellular adhesions, such as VE-cadherin recycling and clustering that regulate intercellular adhesion (Cao et al., 2019). We thus used fluorescence photobleaching after recovery (FRAP) measurements to determine whether observed changes in actin and energy depletion also results in local changes at junctions. Here, we tested whether PFK15 alters VE-cadherin dynamics. Images in Fig. 6a&b show the fluorescence reduction and recovery in control and PFK15 treated cells, respectively. At t=0, the images show the fluorescence reduction, after illuminating a 17 μm^2^ region of the junction.

**Figure 6:**
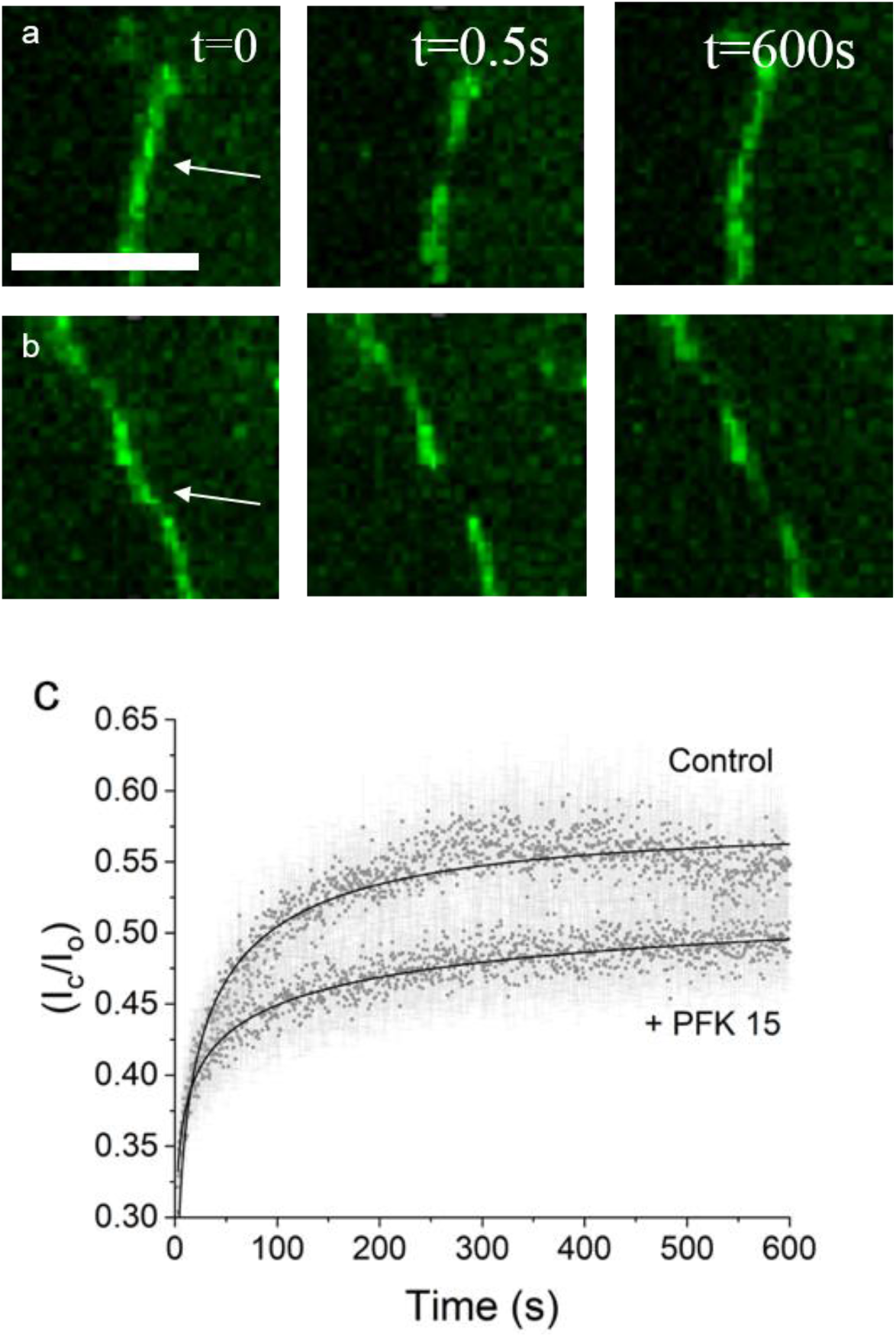
PFK15 treatment alters the recovery time and mobile fraction of VE-cadherin at intercellular junctions. Fluorescent images of GFP-VE-cadherin at junctions between HLMVECs in a confluent monolayer on fibronectin-coated, 40kPa gels. The arrows in a) and b) indicate the region of interest prior to bleaching. a) Control and b) PFK15 treated HLMVECs prior to photobleaching. The bleached region is shown for control and PFK15 treated cells initially and after 10 min. c) Weighted, nonlinear least squares fits of the fluorescence recovery data to equation 1. The best fit parameters are given in the text. (P_Recovery Time_ < 0.001, P_Mobile Fraction_ < 0.01, N_C_ = 7, N_I_ = 8). Scale bar = 25 μm.

Figure 6c shows the normalized intensity versus time, after the initial bleach. The data were fitted to an intensity recovery model that assumes anomalous diffusion (Fig. 6c). Data were fit to the following equation using non linear least squares regression:

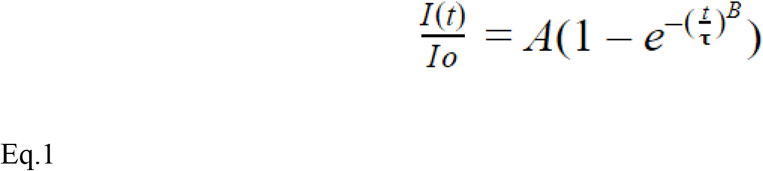

Here, A is the mobile fraction and **τ** is the recovery time (Lorén et al., 2015). The exponent, B indicates anomalous sub diffusion when B<1 and Brownian diffusion when B =1.

The fits in Fig. 6c show that Eq. 1 describes data obtained under both conditions, and B < 1 in both cases. With untreated cells, the fitted parameters for the recovery time and mobile fraction were, respectively 10.9 s ± 0.2 and 0.576 ± 0.002. PFK15 treatment altered both the recovery time and the mobile fraction, with the best-fit parameters being, respectively, 4.7 s ± 0.5 and 0.550 ± 0.008. The exponent B decreased from 0.333 ± .005 to 0.172 ± 0.007. The fluorescence recovery time for the control cells was significantly longer than for PFK15 treated cells (p < 0.001), and the mobile fraction was significantly lower in inhibitor-treated cells (p < 0.01). These results show that PFK15 treatment also perturbs local VE-cadherin dynamics.

## Discussion

Loss of vascular barrier integrity is observed in multiple diseases such as acute respiratory syndrome (ARDS) or diabetic retinopathy, resulting in the massive influx of proteinaceous fluid into the tissue that compromise organ function (Komarova Y, Circ Res 2017). Much of the research on the mechanisms of endothelial junctional integrity and traction forces has focused on the expression levels of the molecular mediators such as VE-cadherin (Komarova Y, Circ Res 2017), but little is known about the metabolic regulation of junctional forces. The main finding of this study is that impairing glycolysis—the principle source of ATP in endothelial cells (De Bock et al., 2013) —reduces tension at cell-cell junctions, in part, by disrupting actin organization and focal adhesions distal from the cadherin-mediated intercellular junctions. Measurements at the macroscopic level of intercellular contacts and at the level of VE-cadherin proteins revealed that inhibiting the glycolysis regulatory enzyme PFKFB3 resulted in a significant drop in junctional tension. Hypoxic conditions are known to potentiate endothelial barrier disruption (Tojo et al., 2018), but most studies have focused on identifying local biochemical changes that destabilize cell-cell contacts. These results reveal that ATP depletion results in global changes that also impinge on intercellular adhesions (Conradi et al., 2017).

The force balance measurements highlight two parameters that contribute to the reduced force on inter endothelial contacts; the shift in the magnitude of the traction stresses and the loss of orientational coherence of traction vectors relative to the junction normal. The latter is visualized using wind rose plots, which reveal a more random distribution in vector angles relative to the junction normal, following PFK15 treatment. Either change individually would reduce the tension, but PFK15 treatment alters both. Importantly, the change in traction stresses used to determine the junction tension was not simply due to the disruption of intercellular adhesions. Cells remained in contact throughout the measurements. Additionally, in isolated cells, PFKFB3 inhibition similarly disrupted the radial orientation of traction force vectors and disrupted actin stress fibers that typically span the cell.

The decrease in the force on cell junctions correlates with the lower tensile force on VE-cadherin complexes, which influence endothelial barrier integrity (Komarova et al., 2017). This might be expected. However, in shear sensing, the onset of fluid flow over endothelial monolayers results in a decrease in force on VE-cadherin and an increase in force on platelet endothelial cell adhesion molecule one (PECAM-1) at junctions (Conway et al., 2013). Here we attribute the lower tension on VE-cadherin, in part, to the disruption of actin fibers to which cadherin is mechanically linked. We did not detect an obvious decrease in VE-cadherin at junctions—a change that would likely increase the force on the remaining cadherin bonds. At the same time, lower tension could destabilize VE-cadherin complexes, by decreasing catenin dependent connections to actin (Buckley et al., 2014, Tabdili et al 2011, Le Duc et al., 2010). Prior studies reported causal relationships between force on cadherin complexes and junction stability (Giannotta et al., 2013). These results show that metabolic dysfunction impacts both protein level mechanics and the overall junction tension, by perturbing the actomyosin cytoskeleton and tethering forces in the cell.

Energy depletion induces actin disintegration and gap formation in endothelial monolayers (Kuhne et al., 1993). The immunofluorescence images and F-actin analyses confirm microfilament disruption, but they also highlight broader impacts on global and junctional mechanics. Endothelial junctions are regulated by a balance of tethering and contractile forces that are propagated through the cytoskeleton (Dudek et al., 2001). In single cells, myosin II and actin networks spanning the cell mechanically couple distal focal adhesions (Cai et al., 2010). Here, immunofluorescence images show that cell-cell contacts are organizing centers that orient actin fibers and traction stresses, which in turn contribute to the force on intercellular adhesions. Actin disintegration severs this connectivity, and thereby decreases the junction tension. At the same time, energy depletion and actin disruption would also reduce contractile forces required to maintain focal adhesions (Bershadsky et al., 2003), accounting for the observed decrease in punctate paxillin staining. Some focal adhesions and actin fibers remain after PFK15 treatment, but the traction generation and mechanical connectivity with cell-cell contacts are significantly impaired.

Tension influences the stability of inter endothelial junctions, but local, ATP-dependent processes such as actin remodeling, as well as cadherin endocytosis, exocytosis and clustering also maintain junctional homeostasis (Wu et al., 2019). PFKFB3 inhibition reduced the fluorescence recovery time (increased mobility), possibly by altering cytoskeletal interactions. Yet, the mobile fraction of VE-cadherin is lower. Although determining the biochemical mechanisms underlying the FRAP results is beyond the scope of this work, the findings confirmed that energy depletion also impinges on local processes at cell junctions. Our findings thus demonstrate that inhibiting glycolysis impacts both local and global processes that regulate adherens junctions.

## Conclusions

Energy depletion destabilizes endothelial adhesions, resulting in gap formation and compromised barrier function. The findings reported here show that inhibition of glycolysis not only impacts the tensile force on VE-cadherin complexes but also disrupts the global actin network and distal focal adhesions that regulate global cell mechanics and the tension at cell-cell junctions. Thus, the impact on endothelial junctions due to impaired glycolysis is not limited to compartments near cell-cell contacts, but also involves global changes in actin and focal adhesions distal from junctions. Understanding this link between glycolysis and junction mechanics highlights the importance of endothelial metabolism in maintaining barriers function during homeostasis. It also suggests that augmenting glycolytic ATP production could be important for restoring junctional integrity and forces during pathological processes.

## Author Contribution

GS designed experiments, performed experiments, wrote TFIMB software, analyzed data, and wrote the manuscript. PG engineered adenovirus constructs for VE-cadTS expression, assisted with lentivirus infections, and wrote the manuscript. PG supplied the adenovirus, PFK15, and HLMVECs. JR designed experiments and edited the manuscript. DL designed experiments, analyzed data, wrote and edited the manuscript.

## Acknowledgements

We thank Hyunjoon Kong and Paul Kenis (University of Illinois at Urbana Champaign) for use of their plasma cleaners. We thank Austin Cyphersmith (Research Specialist at the Institute of Genomic Biology) for assistance with FRAP, FRET, and immunofluorescence imaging. This work was supported by AHA 18PRE34070092 to PG, and PO1 HL060678-16 to DEL and JR.

## Supplementary

**Figure S1:**
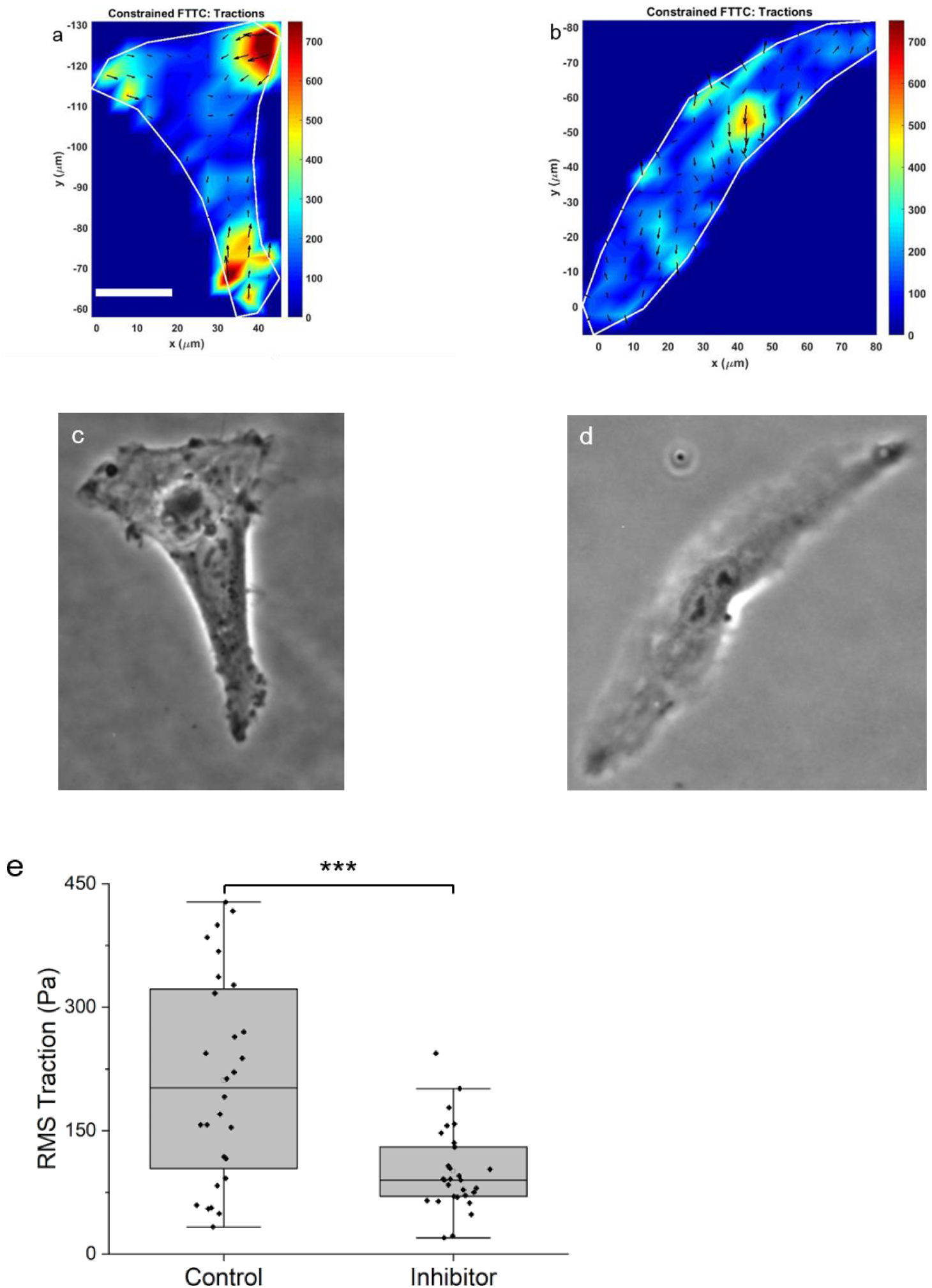
PFK15 treatment reduces the traction force amplitude and orientational uniformity. a) Traction force heat maps of treated and b) PFK15 treated, single HLMVECs. Cells were seeded at low density on fibronectin coated 40kPa gels. DIC images of c) control and PFK15 treated HLMVECs. e) Root mean square traction stress (Pa) generated by HLMVECs 40 min after PFK15 treatment, compared with untreated controls. Cells were seeded on the substrates described in parts a and b (p < 0.001, N_C_ = 28, N_I_ = 29). Scale bar is 20 μm.

**Figure S2:**
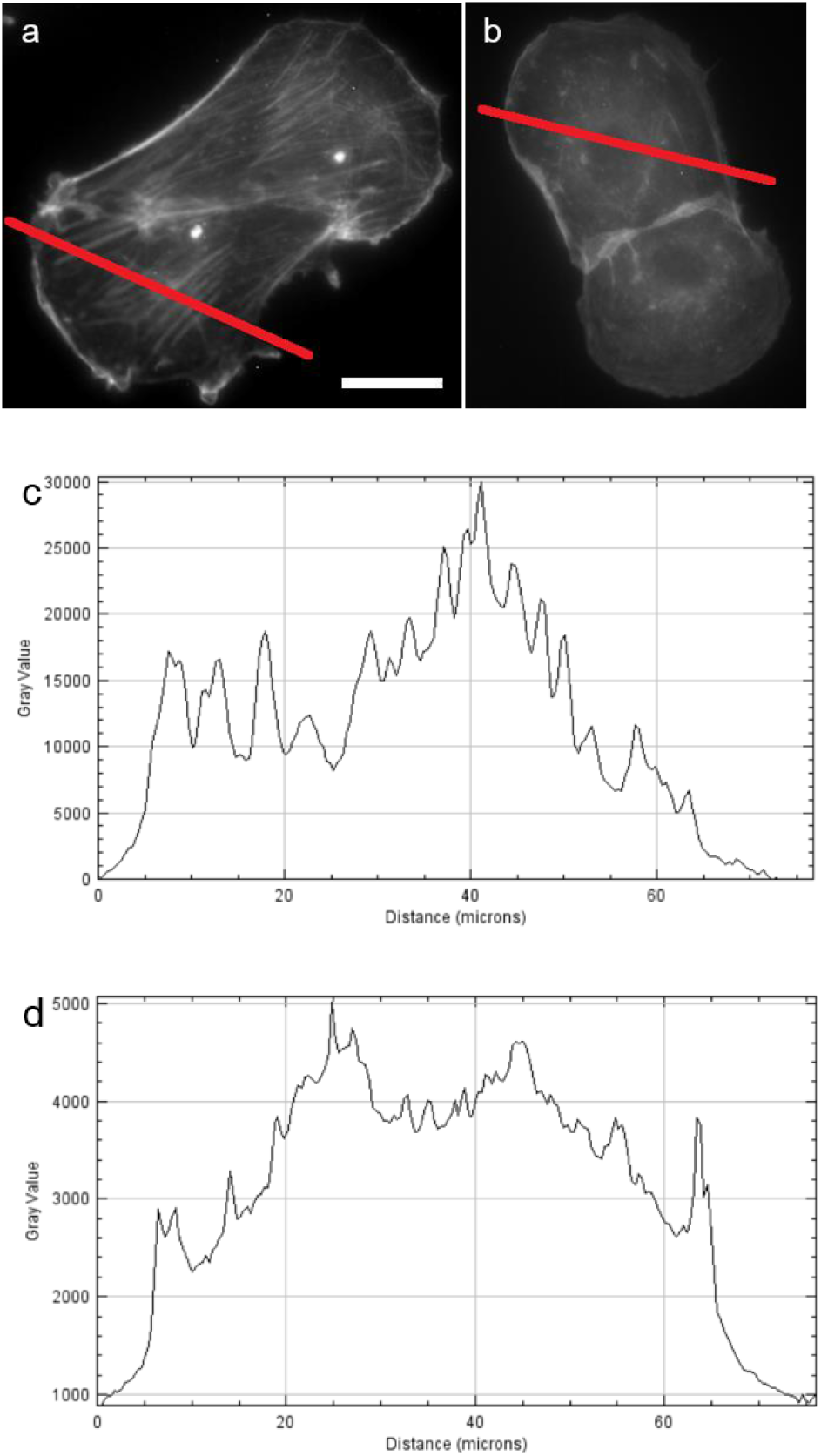
PFK15 alters the width and length of F-actin fibers. Immunofluorescence images of F-actin in HLMVECs in a) controls and b) after PFK15 treatment. The red line indicates the showing where intensity scans were taken. c) Intensity scans across the cell in panel a and d) intensity scan across the cell in panel b. F-actin intensity scans in HLMVECs show distinct peaks corresponding to F-actin fibers whereas the actin is more diffuse and the fibers are narrower and shorter in PFK15 treated cells. Cells were seeded on fibronectin patterns on 40 kpa gels. Scale bar = 25 μm.

**Figure S3:**
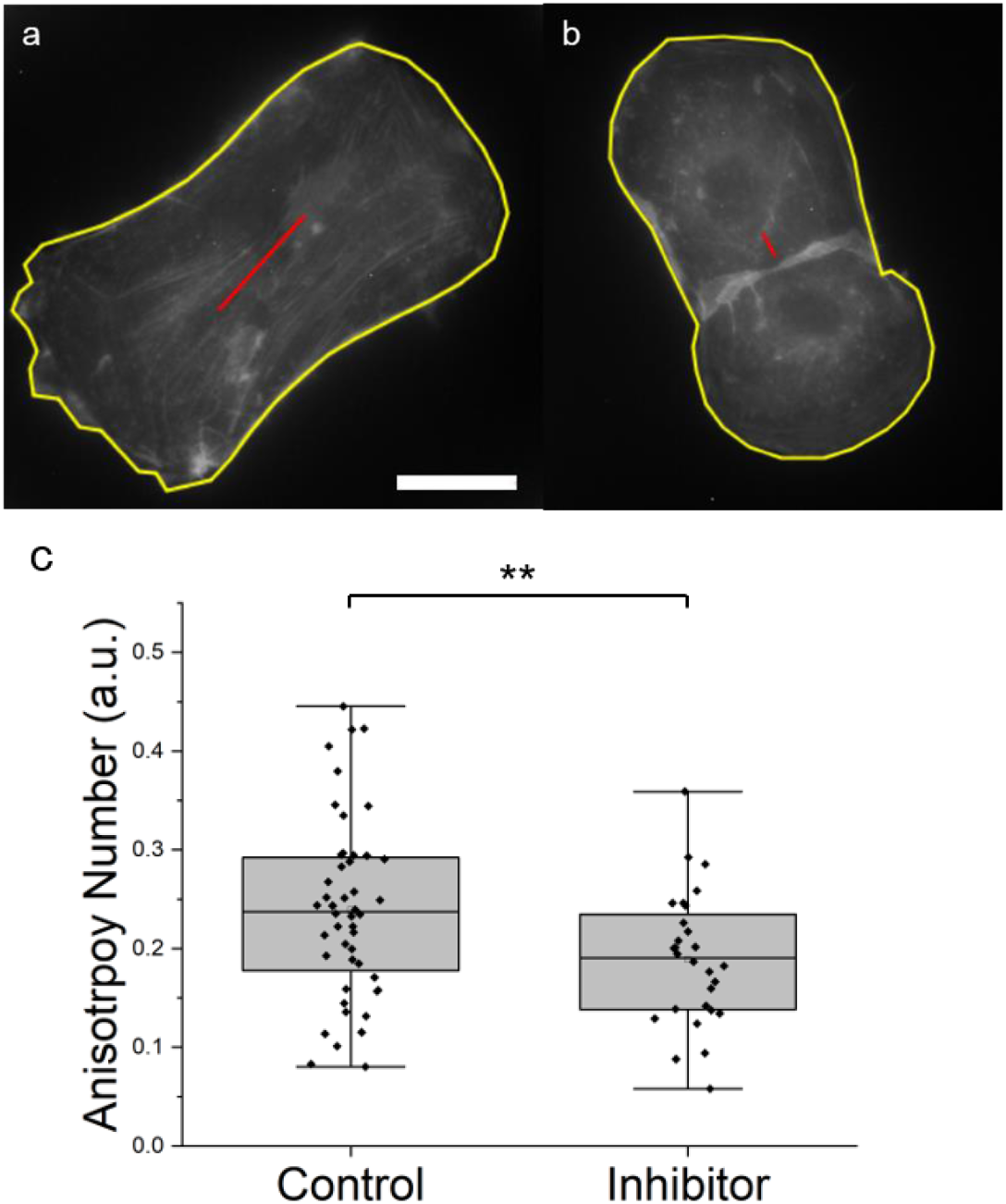
Anisotropy of F-actin fibers decreases following PFK15 treatment. The anisotropy orientation and magnitude corresponding to F-actin fibers within the cell boundary (yellow outline). Cells were seeded on microcontact printed fibronectin patterns on 40 kPa gels. Phalloidin-stained actin was imaged at 60x magnification. Quantified actin anisotropy is shown for a) untreated controls and for b) PFK15 treated HLMVECs (generated by Fibriltool). c) Average anisotropy number of actin fibers in untreated and treated cells. The difference is statistically significant (**p < 0.002, N_C_ = 46, N_I_ = 33). Scale bar = 25 μm.

